# Evidence for the equilibrium between monomers and dimers of the death domain of the p75 neurotrophin receptor

**DOI:** 10.1101/2021.05.02.442373

**Authors:** Zhen Li, Zhi Lin, Carlos F. Ibáñez

## Abstract

The p75 neurotrophin receptor (p75^NTR^) is an important mediator of synaptic depression and neuronal cell death, and its expression increases upon nerve injury and in neurodegenerative diseases. However, the molecular mechanisms leading to the activation of this receptor are still a matter of debate. The oligomerization properties of the death domain (DD) of p75^NTR^ are critical for our understanding of the activation mechanisms of the receptor. In this paper, we present additional evidence supporting the existence of an equilibrium between monomeric and dimeric forms of the p75^NTR^ DD in solution and in the absence of any other protein. Dynamic light scattering (DLS) measurements of native, untagged human p75^NTR^ DD at room temperature yielded Rh=2.11 for this domain in 20mM phosphate buffer, corresponding to a molecular weight (MW) of approximately 19kDa, much closer to the theoretical MW of the homodimer (i.e. 21kDa) than the monomer. MWs deduced from the Rh of different control proteins used as standards were all congruent with their theoretical MWs. In addition, size-exclusion FPLC profiles of un-tagged human p75^NTR^ DD in both HEPES and phosphate buffers revealed elution volumes corresponding to a MW of about 15kDa, which is intermediate between monomer and dimer, and indicative of dynamic monomer/dimer interconversion during the run. Together with our previous NMR studies, as well as biophysical data for other investigators, these results support the notion that the DD of p75^NTR^ exists in equilibrium between monomers and dimers in solution, a notion that is in agreement with the oligomerization properties of all members of the DD superfamily.

## Introduction

The p75 neurotrophin receptor (p75^NTR^, also known as NGFR and TNFRSF16) can induce neuronal and glial cell damage, axonal degeneration and synaptic dysfunction (Ibáñez and Simi, 2012). p75^NTR^ is a central piece of several neurodegeneration pathways, including nerve injury and Alzheimer’s Disease. Understanding how this receptor becomes activated and signals in response to neural damage is critical to efforts aimed at targeting p75^NTR^ signaling for the development of therapeutic strategies. The cytoplasmic domain of p75^NTR^ contains a C-terminal globular protein module of 80 amino acid residues with a characteristic 6-helix bundle fold known as the death domain (DD) (Liepinsh et al., 1997). DD-containing proteins play central roles in apoptotic and inflammatory signaling through the formation of oligomeric protein complexes. Fluorescence resonance energy transfer (FRET) experiments have shown that the two DDs in the p75^NTR^ receptor dimer are in close proximity to each other (high FRET state) and that NGF binding induces a decrease in FRET signal (Sykes et al., 2012; Vilar et al., 2009). In agreement with the FRET results, solution NMR studies from different laboratories, including ours, have demonstrated that the p75^NTR^ DDs exists in equilibrium between monomeric and dimeric forms depending on pH and counter ion (Lin et al., 2015; Vilar et al., 2014). The apparent Kd of dimerization derived from homo-FRET anisotropy experiments was 49±15 µM (Lin et al., 2015). This relatively low-affinity interaction may facilitate DD separation (low FRET state) upon receptor activation by neurotrophins. Ligand-induced separation of p75^NTR^ DDs allows the recruitment of intracellular components for downstream signal propagation. In a recent article, Goncharuk et al. (Goncharuk et al., 2020) challenged evidence on the intrinsic the oligomerization properties of the p75^NTR^ DD. They argued that the p75^NTR^ DD is monomeric in solution and propose that formation of DD dimers within cells requires the aid of a yet-to-be-identified “helper” protein.

At the heart of this issue, is the notion that p75^NTR^ itself exists as preformed homodimers in the plasma membrane of neurons and other cells in its native state (a fact not disputed by Goncharuk et al.) and that this drives an equilibrium between monomeric and dimeric states of intracellular DDs. Our solution NMR structure of the DD homodimer (Lin et al., 2015) showed that critical residues mediating interaction of p75^NTR^ with downstream effector molecules, such as RIP2, are buried in the DD dimer interface, indicating that p75^NTR^ activation entails conformational changes that separate the DDs to allow binding of intracellular molecules for downstream signal propagation. Thus, the oligomerization properties of the DD in p75^NTR^ are critical for our understanding of the activation mechanisms of the receptor and our ability to design strategies that may interrupt it. In this communication, we present additional evidence supporting the equilibrium between monomeric and dimeric forms of this domain in solution and in the absence of any other protein.

## Results and Discussion

The notion that the DDs in p75^NTR^ dimers are in close proximity of each other, and may form homodimeric complexes, was first advanced in homo FRET studies showing an intrinsically low anisotropic state (i.e. high FRET) of intracellular domains of p75^NTR^ at resting conditions (Vilar et al., 2009). Ligand activation resulted in decreased FRET (increased anisotropy), suggesting separation of intracellular domains. Other laboratories independently obtained similar results (Sykes et al., 2012). Importantly, these observations were made in very different cell types, neuronal and non-neuronal, of different embryonic origins and animal species, many of which do not normally express p75^NTR^; a fact that does not easily lends itself to support the existence of a specific “helper” protein aiding DD dimerization, as proposed by Goncharuk et al. From our earlier paper (Vilar et al., 2009), we also know that eGFP-tagged DDs of p75^NTR^ show concentration-dependent anisotropic changes consistent with protein homodimerization in solution, with a Kd of ≈ 50μM (Figure 5—figure supplement 1 E in ref. (Lin et al., 2015)), also supporting a steady-state equilibrium between monomers and dimers in solution. We also note that Goncharuk et al. did not mention the intermolecular NOEs that were detected in our sample of ^13^C,^15^N-labeled DD mixed with unlabeled DD (Figure 5—figure supplement 1B in ref. (Vilar et al., 2009)). These NOEs are direct evidence of DD homodimerization and cannot be interpreted in any other way.

The theoretical molecular weight (MW) of the protomer of p75^NTR^ DD is 10.5kDa. Our original dynamic light scattering (DLS) measurements of His-tagged human p75^NTR^ DD at ≈ 0.2mM in phosphate buffer showed a hydrodynamic radius (Rh) of 2.06±0.05 nm, corresponding to a ≈ 18 kDa globular protein. We note that the His tag here contributes with 1.8kDa. In HEPES buffer, a Rh of 1.65±0.04 nm (∼10 kDa) was obtained (Figure 5—figure supplement 1C and D in ref. (Vilar et al., 2009)). This result indicates that the DDs in p75^NTR^ can equilibrate between monomeric and dimeric states at relatively low concentration in phosphate buffer solution. Intriguingly, Goncharuk et al. own DLS measurements yielded a Rh of 1.8nm (Goncharuk et al., 2020), corresponding to a protein of ≈ 14kDa, a result that is not compatible with a purely monomeric species. We note that the method used by these authors to measure Rh and derive MW from DLS was not described in their original paper. We have recently repeated our original DLS study using native, untagged p75^NTR^ DD at room temperature, alongside different control monomeric proteins to serve as molecular standards. Representative Rh distributions of the p75^NTR^ DD and the three control samples are shown in Fig. 1A; average Rhs are shown in Fig. 1B. These measurements yielded Rh=2.11 for the untagged human p75^NTR^ DD in 20mM phosphate buffer, corresponding to a MW of approximately 19kDa, much closer to the theoretical MW of the homodimer than the monomer of this domain. We note that the MWs deduced from the Rh of the different control proteins used as standards were all congruent with their theoretical MWs, indicating that our DLS measurements are indeed accurate and reproducible. We have also run size-exclusion FPLC profiles of un-tagged human p75^NTR^ DD in HEPES and PO4 buffers. This analysis revealed elution volumes corresponding to a MW of about 15kDa (Fig. 2), which is intermediate between monomer and dimer, and indicative of dynamic monomer/dimer interconversion during the run. Together with our previous NMR studies, including the solution structure of the DD homodimer, these DLS and FPLC results support the notion that the DD of p75^NTR^ exists in equilibrium between monomers and dimers in solution.

**Figure 1.**
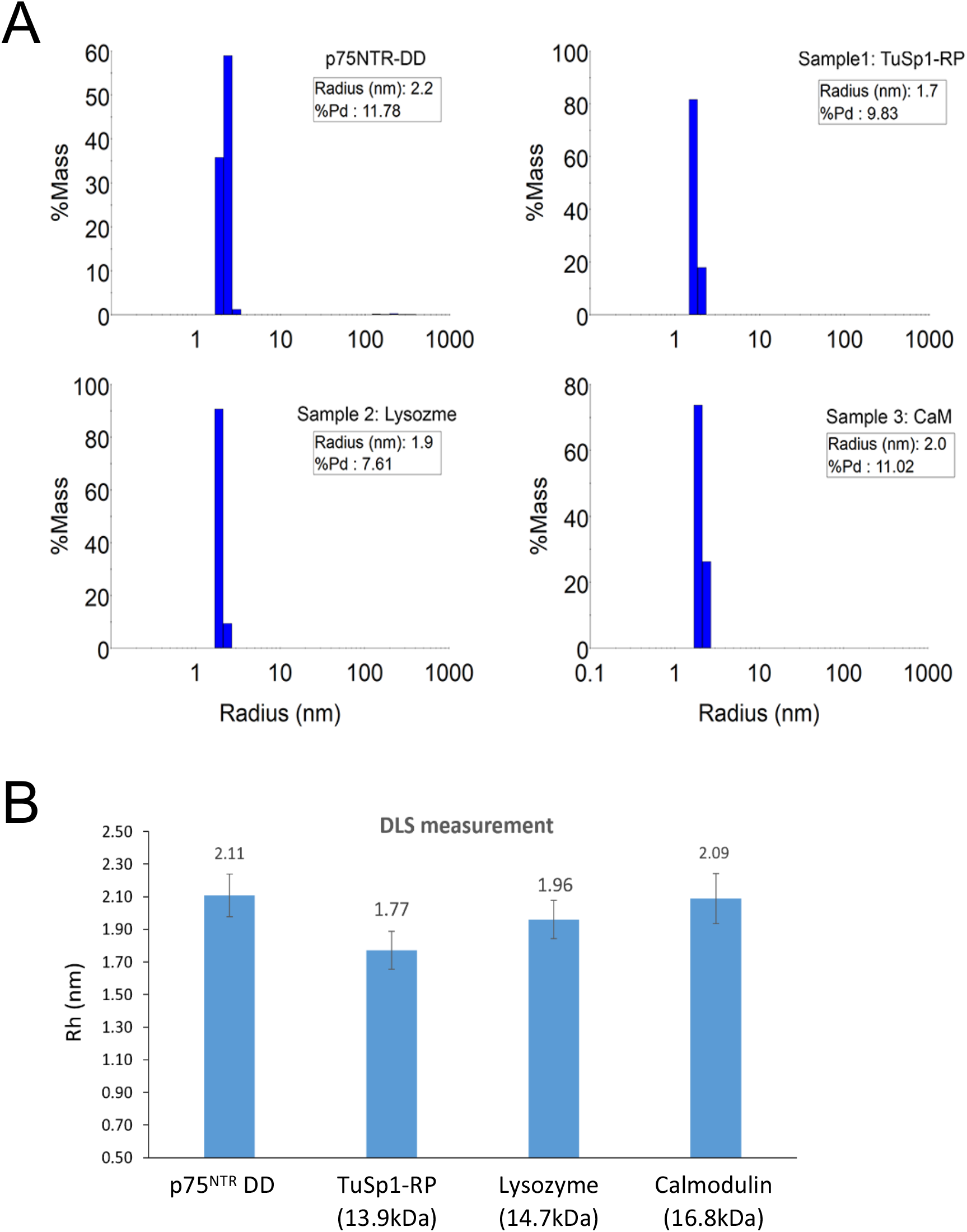
DLS measurements of untagged human p75^NTR^ DD and three control proteins at room temperature. (A) Representative Rh distribution of the p75^NTR^ DD (theoretical protomer MW=10.5kDa) at 3mg/ml in 20mM PO4 buffer, 1mM DTT, pH7.5, and three control samples: 1) His-tagged TuSp1-RP (MW=13.9kDa) at 1mg/ml in 20mM Tris, pH7.5; 2) lysozyme (SangonBiotech, MW=14.7kDa) at 1mg/ml in 20mM PBS; and 3) calmodulin without Ca^2+^ (CaM, MW=16.8kDa) at 1mg/ml in 20mM HEPES, 50mM EGTA, ph7.0 (B) Average Rh of p75^NTR^ DD and the three control samples.

**Figure 2.**
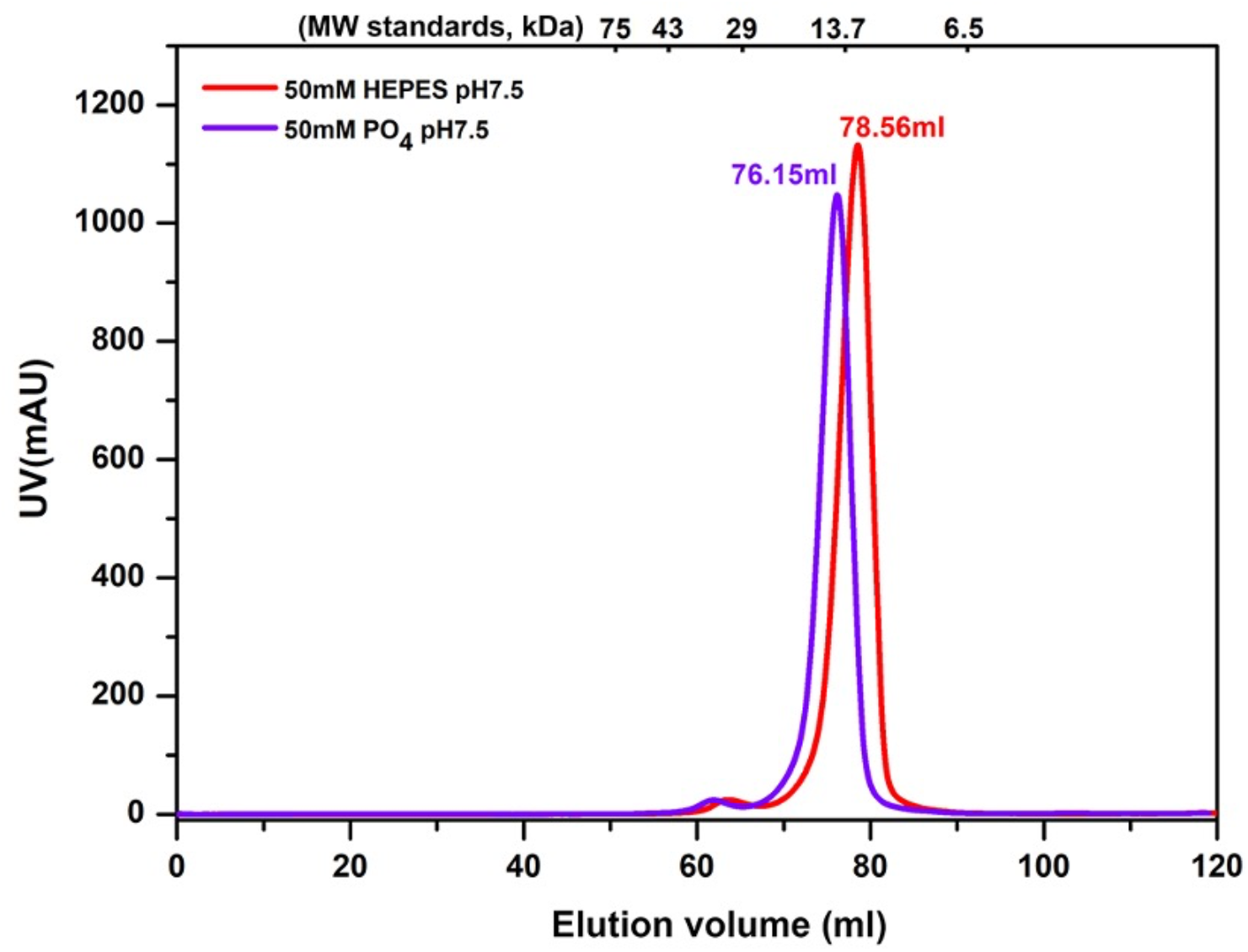
FPLC profiles of human p75^NTR^ DD in HEPES and phosphate buffers. FPLC profiles (Superdex 75) of human p75^NTR^ DD in HEPES (red) and phosphate (blue) buffers at pH 7.5. MW standards from 6.5 to 75 kDa are indicated above the profiles. Elution volume at 76.15ml corresponds to a MW of ∼15kDa, in agreement with the notion that p75^NTR^ DDs equilibrate between monomers and dimers in solution and in different buffers.

Biophysical data from other investigators also support the existence of dimers of the DD of p75^NTR^ in solution. As shown by Vilar et al. (Vilar et al., 2014), analytical ultracentrifugation data on the intracellular domain of p75^NTR^ (which includes the DD and a 6.5 kDa juxtamembrane domain) in PBS buffer demonstrated that this protein behaves in solution as a single species with MW 30.7 kDa, close to the theoretical dimer of the p75^NTR^ intracellular domain (Figure 3 in ref. (Vilar et al., 2014)). Also, in the same paper, Vilar et al. reported concentration-dependent chemical shift changes of ^15^N-labeled p75^NTR^ intracellular domain that were indicative of self-assembly with a Kd of ≈ 100μM. Importantly, mutation of critical residues in the dimerization interface significantly disrupted dimer formation (Figure 4 in ref. (Vilar et al., 2014)).

In summary, these observations strongly indicate that monomeric and dimeric states of the DD of p75^NTR^ exist in equilibrium at physiological conditions. This supports an intrinsic ability of this protein to self-associate in solution, in congruence with the oligomerization properties of all members of the DD superfamily. This is also a more parsimonious explanation than speculations on the existence of an unknown “helper” protein mediating p75^NTR^ dimerization universally present in all cell types.

## Methods

The cDNA of human p75^NTR^ DD (330-427, 10.5kDa) was subcloned into a pET-32-derived expression vector. The His-tagged p75^NTR^ DD was expressed in SoluBL21 (DE3) in LB medium and purified by using Ni-NTA affinity chromatography. The His tag at the N-terminal was cleaved by tobacco etch virus (TEV) protease and the sample was applied to the second Ni-NTA affinity chromatography to remove the His tag. FPLC gel filtration (HiLoad 16/600 Superdex 75, ÄKTA), and/or ionic exchange (MonoQ, AKTA) were further used to purify the untagged p75^NTR^ DD. The hydrodynamic radii (Rh) of the untagged p75^NTR^ DD and three control proteins used for MW calibration were measured by dynamic light scattering (DLS, DynaPro) at 20°C. The DLS data were analyzed by using Dynamics 7.8 software. All the experiments were done in the same machine and temperature conditions.

## Author contributions

Zhen Li generated the data, Zhi Lin designed and supervised the research, and commented on the original manuscript, C.F. Ibáñez wrote the manuscript.

## Additional Information

The authors declare that they have no competing interests.

